# AlGrow: a graphical interface for easy, fast and accurate area and growth analysis of heterogeneously colored targets

**DOI:** 10.1101/2024.03.04.583395

**Authors:** Marcus McHale, Ronan Sulpice

## Abstract

Image analysis is widely used in plant biology to determine growth rates and other phenotypic characters, with segmentation into foreground and background being a primary challenge. Statistical clustering and learning approaches can reduce the need for user input into this process, though these are computationally demanding, can generalise poorly and are not intuitive to end users. As such, simple strategies that rely on the definition of a range of target colors are still frequently adopted. These are limited by the geometries in color space that are implicit to their definition; i.e. thresholds define cuboid volumes and selected colors with a radius define spheroid volumes. A more comprehensive specification of target color is a hull, in color space, enclosing the set of colors in the image foreground. We developed AlGrow, a software tool that allows users to easily define hulls by clicking on the source image or a three-dimensional projection of its colors. We implemented convex hulls and then alpha-hulls, i.e. a limit applied to hull edge length, to support concave surfaces and disjoint color volumes. AlGrow also provides automated annotation by detecting internal circular markers, such as pot margins, and applies relative indexes to support movement. Analysis of publicly available Arabidopsis image series and metadata demonstrated effective automated annotation and mean Dice coefficients of >0.95 following training on only the first and last images in each series. AlGrow provides both graphical and command line interfaces and is released free and open-source with compiled binaries for the major operating systems.

## BACKGROUND

In plant and macroalgal species, non-destructive estimates of growth rate and other phenotypic characters can be determined from 2-dimensional (2D) images captured with digital cameras (Leister et al., 1999; Fort et al., 2019). Although 3-dimensional (3D) scanning and other depth measuring tools such as time-of-flight cameras may improve accuracy, the hardware for such techniques is more expensive and less accessible (Mccormick et al., 2016). Further, multiple viewing angles may be used to reconstruct 3D models and improve accuracy in species with complex architecture, reinforcing the role of digital cameras in plant phenotyping (Cabrera-Bosquet et al., 2016). Outside of plant phenotyping, similar imaging techniques may be used in classification for automated weed management in the agricultural sector, though variable field conditions present additional challenges in this context (Hamuda et al., 2016).

Typical digital cameras capture trichromatic RGB values (Red, Green and Blue) to represent color for each pixel. The Commission Internationale de l’éclairage (CIE) designed the earliest and most widely used such encoding, CIE RGB, from earlier experiments with human observers (Smith and Guild, 1931). The CIE later designed the CIELAB color space, a transformation of RGB, such that Euclidean distance is approximately proportional to perceived color difference under similar luminance (The Optical Society, 1948). The color dimensions in CIELAB are termed L*, a* and b* (asterisks are used to distinguish from a similar earlier definition), representing “lightness to darkness”, “redness to greenness” and “blueness to yellowness”, respectively, (The Optical Society, 1948). Distance in CIELAB is termed delta E (ΔE), after “*Empfindung”* in German, which translates to “sensation” in English.

A great diversity of software tools have been developed to support RGB image analysis in plant phenotyping for research, varying from software libraries for customized pipeline development, to complete software packages (Lobet et al., 2013; Gehan et al., 2017; Minervini et al., 2017; Henke et al., 2021). Segmentation and classification into target and background using such software is most simply achieved by applying fixed or dynamic thresholds to the color dimensions. This is either performed in the native color space (RGB), a transformed representation (e.g. CIELAB), or across multiple such representations (Leister et al., 1999; Fort et al., 2019). Part of the benefit of thresholds in image segmentation is the ease of implementation of a simple user interface for their determination, for example, the sliders implemented in ImageJ (Schneider et al., 2012). However, applying thresholds is limited to cuboid boundaries in a 3D color space, with more complex selections requiring iterative selections.

Another strategy, employed in CoverageTool, intuitively supports the selection of a set of target colors from the source image, each with a color distance radius (termed tolerance level) applied as a threshold to classification for both foreground and background (Merchuk-Ovnat et al., 2019). This strategy is however similarly limited to spheroid boundaries, with more complex selections requiring mor sampled colors, each with smaller radii. CoverageTool is currently limited to the selection of ten foreground and ten background colors and uses color spaces that do not conform to user perceptions of color differences.

A more recent semi-automated segmentation strategy that still only relies on pixel color information, aggregates the dimensions of multiple color spaces into a set of representative “eigen-colors” by principle/independent component analysis (PCA/ICA) and applies k-means clustering in this hyper-dimensional space followed by manual selection of target clusters (Henke et al., 2021). This strategy reduces the need for user input, however due to the dimensional transformation the resulting selections may not be applied across multiple images. K-means was selected by the authors for it’s performance over other algorithms, though computation time is still approximately 1-minute for a single 5 megapixel (MP) image. K-means also implicitly defines (hyper)-spheroid selections, requires a suitable value for the number of clusters (k) to ensure appropriate fits, and performs poorly where clusters are of differing sizes, densities and shapes.

We previously reported a custom image capture and analysis pipeline for multiplexed arrays of macroalgal lamina discs and applied this to an assessment of *Ulva spp*. diversity (Fort et al., 2019). This pipeline employed the PlantCV framework to initially mask background using fixed thresholds, and subsequently relied on the ImageJ graphical interface to interactively annotate image stacks and apply further thresholds for segmentation (Schneider et al., 2012; Fahlgren et al., 2015). The manual annotation and application of thresholds to images in this pipeline was time-consuming and susceptible to operator error and/or biases. Further, in the continued application of this pipeline to *Ulva spp*., and in extension to other species such as *Palmaria palmata*, we encountered issues in segmenting images with less distinct color boundaries and more varied color distributions.

These difficult to segment images typically represent variation in subject color or illumination, microalgal contamination and/or exudate/leachate from tissues. Shape and textural properties that are frequently considered in plant image segmentation were thought to be unreliable predictors in this context (Minervini et al., 2014). We considered the possibility of improving color specification with convex hulls to define boundaries of a target color space. We then generalized this to support concave surfaces and/or disjoint color volumes with the simple yet flexible definition of an alpha hull, i.e. a limit to the length of edges considered during hull construction (Edelsbrunner et al., 1983).

Here we provide AlGrow, a tool to interactively select hull vertices for color based plant and macroalgal image segmentation. AlGrow also implements an effective automated annotation strategy for multiplexed images and provides detailed plots, debugging figures, regression-based growth rate analyses and reports from the resulting data

## RESULTS

The first steps in AlGrow configuration, defining a scale and selecting a color that highlights circular markers for regions of interest, are each achieved by clicking on the image (Figs. S1 and S2). High contrast circles of a defined range of sizes are then readily detected, for example in images of *Palmaria palmata* from the previously described macroalgal phenotyping apparatus (Fig. 1) (Fort et al., 2019). Hierarchical clustering of detected circles into plates of known sizes is effective in filtering out partially imaged plates and reflections (Fig. 1A). Similar clustering into rows or columns, performed for plates and again within plates, provides consistent annotation that is robust to movement where approximate relative positions are maintained (Fig 1B). Debugging output, such as cladograms of circle center distance, with the determined cut-height, allow users to diagnose issues in layout detection (Fig. S3).

**Figure 1.**
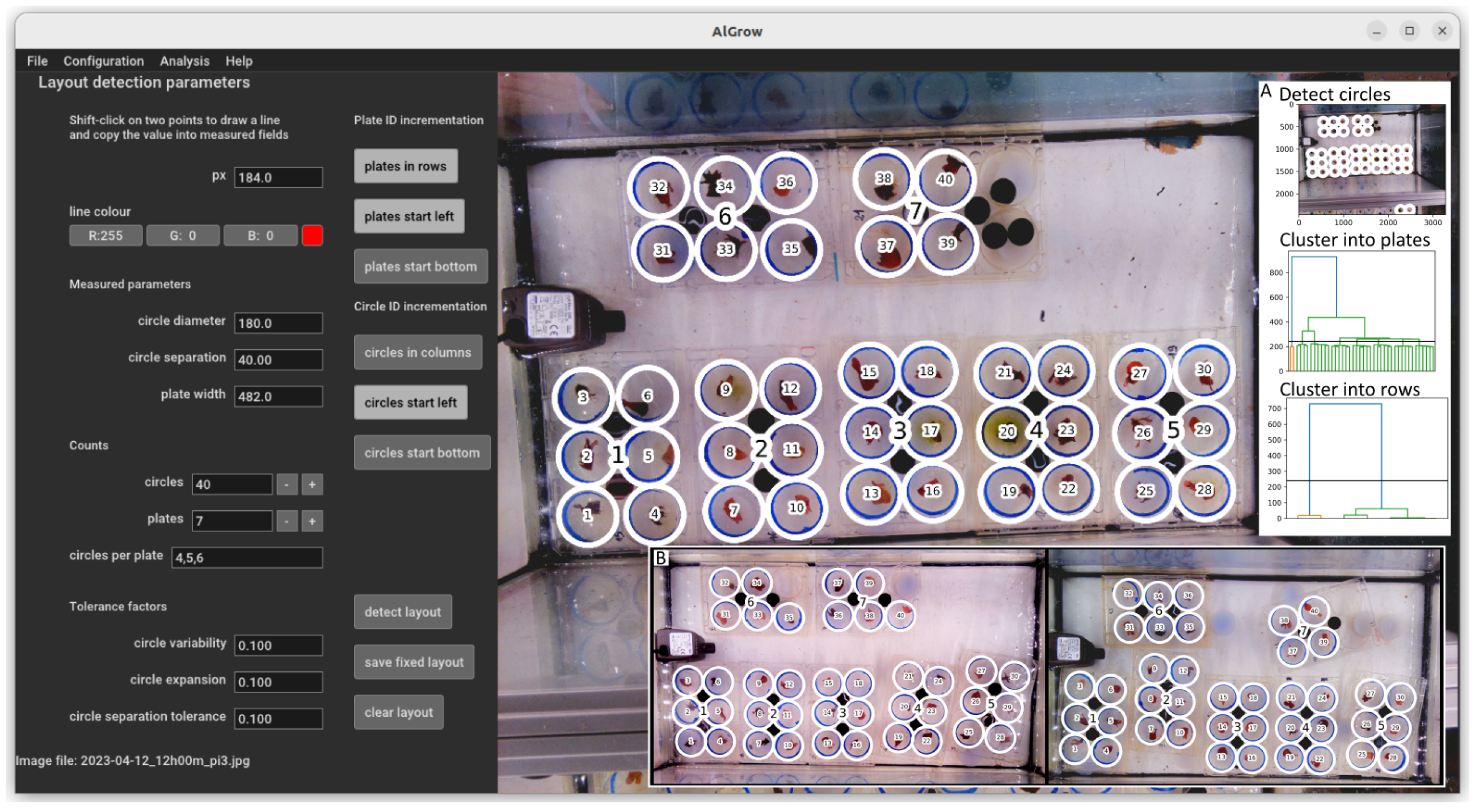
Circle detection, clustering and ID incrementation on an image of *Palmaria palmata (*supplementary file: “palmaria.jpg”). In the AlGrow GUI, a line can be drawn over the loaded image and its span is reported in pixels (px). This facility is used to determine circle diameter, circle separation (the distance between circle edges within a plate) and plate width. Fields are provided to specify counts for the number of circles to detect in the image, the number of plates and the number of circles per plate (multiple options as a comma separated list). Tolerance factors may be supplied to affect circle variability (the range of circle diameters to consider), circle expansion for region of interest specification and circle separation tolerance for plate clustering. Incrementation options are also provided to support alternative ordering of plate and circle IDs. When the “detect layout” button is clicked an attempt to identify the layout is made. When this is successful, an overlay is displayed detailing the detected circles and plates with corresponding IDs. The provided parameters may be used to dynamically detect a layout in each image to support movement across an image series, or alternatively a fixed layout may be saved where movement is not a concern. **Inset A:** Following initial circle detection, clustering removes detected circles that are not within expected clusters. This allows the removal of partially imaged plates. Plates are then clustered into rows (or columns) for annotation, and within each plate similar row or column clustering is applied to establish the order for ID incrementation. **Inset B:** This automated annotations strategy is robust to varying arrangements (left) and/or movement (right).

Target hull specification supports inspection of color distributions as voxels in the 3D CIELAB color space (Fig. 2A). Filtering voxels by the number of pixels they represent allows users to visually identify clusters and determine suitable boundaries for image segmentation (Fig. 2B). Target color boundaries may be defined with as few as four points to form a convex hull. These points may be selected either directly in the image, or from the 3D plot (Fig. 2). The alpha parameter supports more complex topologies and/or disjoint selections by limiting the edge length during hull construction (Fig. S5). The delta parameter specifies the distance (ΔE) from the hull surface to consider as within the target color space. This allows for inaccuracy in the selection of hull vertices and can be used to adjust the sensitivity and specificity of the classification model.

**Figure 2.**
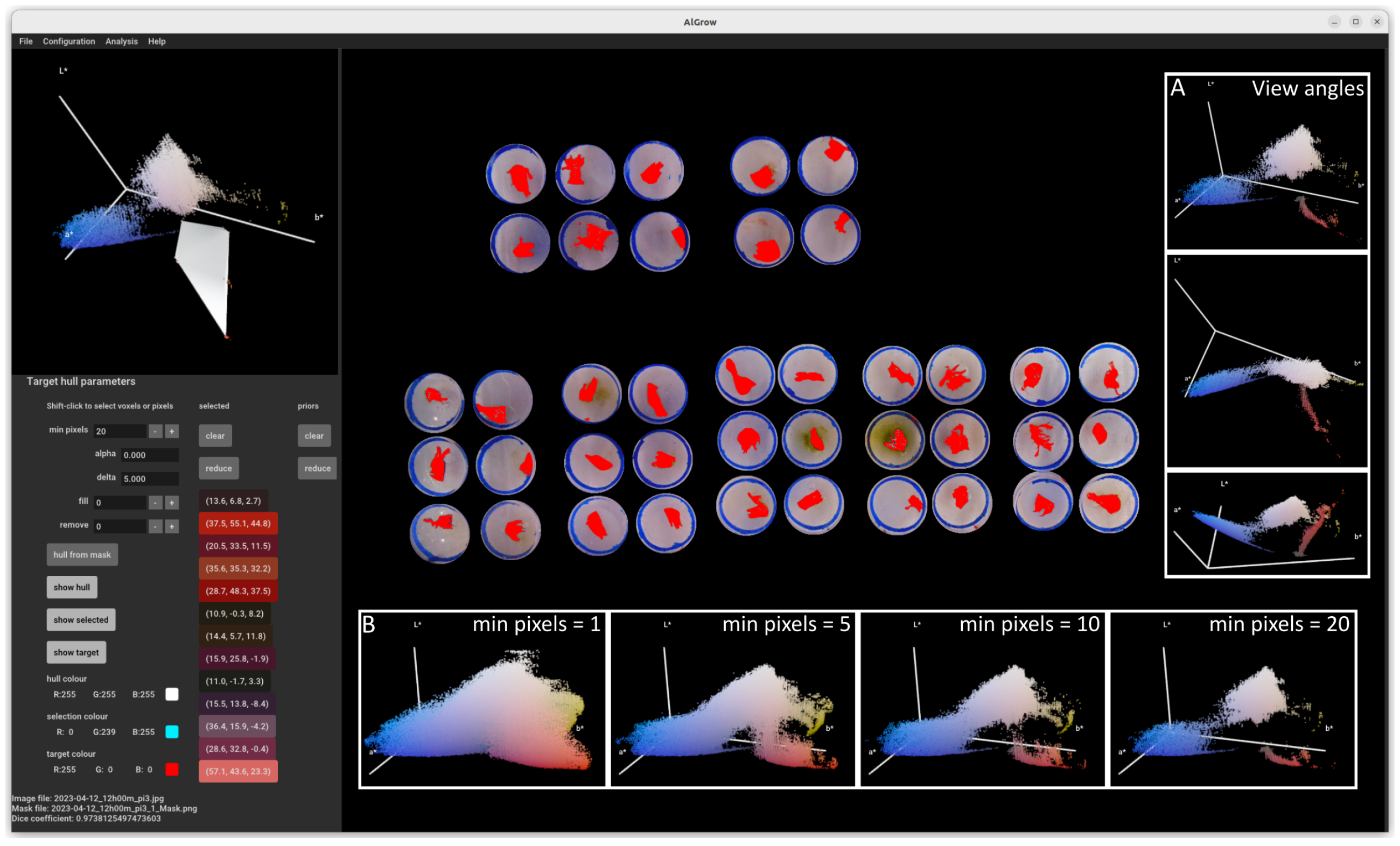
Target hull specification and image segmentation on the image of *Palmaria palmata* (supplementary file: “palmaria.jpg”). Following prior layout detection to mask background in the loaded image (right), voxels representing pixel colors are displayed as points in a 3D plot of the CIELAB color space (top-left, see inset **A** for other viewing angles). Visualization of target clusters is achieved by filtering voxels by the number of pixels that each represents (min pixels = 20, see inset **B** for other values for this parameter). Voxels may be selected/deselected either from the 3D-plot or by clicking the corresponding pixels in the image. Selected voxels are highlighted as spheres in the 3D plot and optionally highlighted in a customizable overlay (cyan) on the image (show selected). When a valid hull is constructed from the selected points at the supplied alpha value (alpha = 0 specifies the convex hull) it is optionally displayed in this plot (show hull). An optional overlay (red) on the target image highlights pixels whose corresponding voxel is either within the hull or within the supplied delta (ΔE = 5) of its surface (show target). Selected voxels are represented by a button labelled with their corresponding coordinates in the “selected” column, where they may be removed individually, cleared, or reduced to a set consisting of only the current hull vertices. To support target hull design across a series of images, when a new image is loaded the current selected voxels are copied to the list of “priors”, also used in hull construction. Priors are similarly used when hull vertices are loaded from a configuration file or command-line arguments. For comparison across methodologies, a target mask may be loaded and the Dice coefficient is reported (0.97, bottom left). This convex hull was generated with less than 1 minute of user interaction, while the loaded mask (supplementary file: “palmaria_alpha_mask.png”) is from a more refined alpha hull generated with approximately 10 minutes of user interaction (Fig. S3). Fill and remove are not used in this figure to most clearly present the accuracy of color based segmentation, however these functions are useful to account for local variation and/or materials that interfere with imaging, such as the nylon mesh in the macroalgal phenotyping apparatus. Configuration for this figure is described in supplementary file “palmaria.conf”.

To compare classification success across different configurations or tools, a foreground mask image can be loaded and the Dice similarity coefficient is reported (Figs. 2, S4 and S5, bottom left). A foreground mask may also be used to select points for hull construction (“hull from mask”). To best represent the effectiveness of the hull in color specification, optional fill and remove functions are not displayed here, though these are effective tools to account for signal noise or other variation such as the nylon mesh in the macroalgal imaging apparatus.

Hull specification may be performed across multiple images to ensure the target color model is sufficiently generalized for an image series. When a new image is loaded, any selected voxels are copied to a list of priors that are also considered as points for hull construction. Points saved in configuration files or provided as arguments are also loaded as priors.

We have demonstrated AlGrow on a public dataset of Arabidopsis images with manually curated foreground masks. We selected the Ara2012 (Raspberry Pi) and Ara2013 (Canon) tray datasets, obtained from https://www.plant-phenotyping.org/datasets-home, as these are also able to demonstrate automated detection of pots (Minervini et al., 2015). A hull trained using only the first and last image of the 16 image Ara2012 series achieved a Dice coefficient of similarity to the manually curated foreground masks of 0.963 without filling small voids or removing small objects and 0.969 with this additional processing (Fig. 3, Table S1). This is comparable to the reported accuracy of other segmentation strategies for these images, including procedures that consider additional texture information (0.964) (Minervini et al., 2014). Segmentation of the Ara2013 dataset achieved similar accuracy, with a Dice coefficient of ∼0.96 across the 27 image series (Table S1).

**Figure 3.**
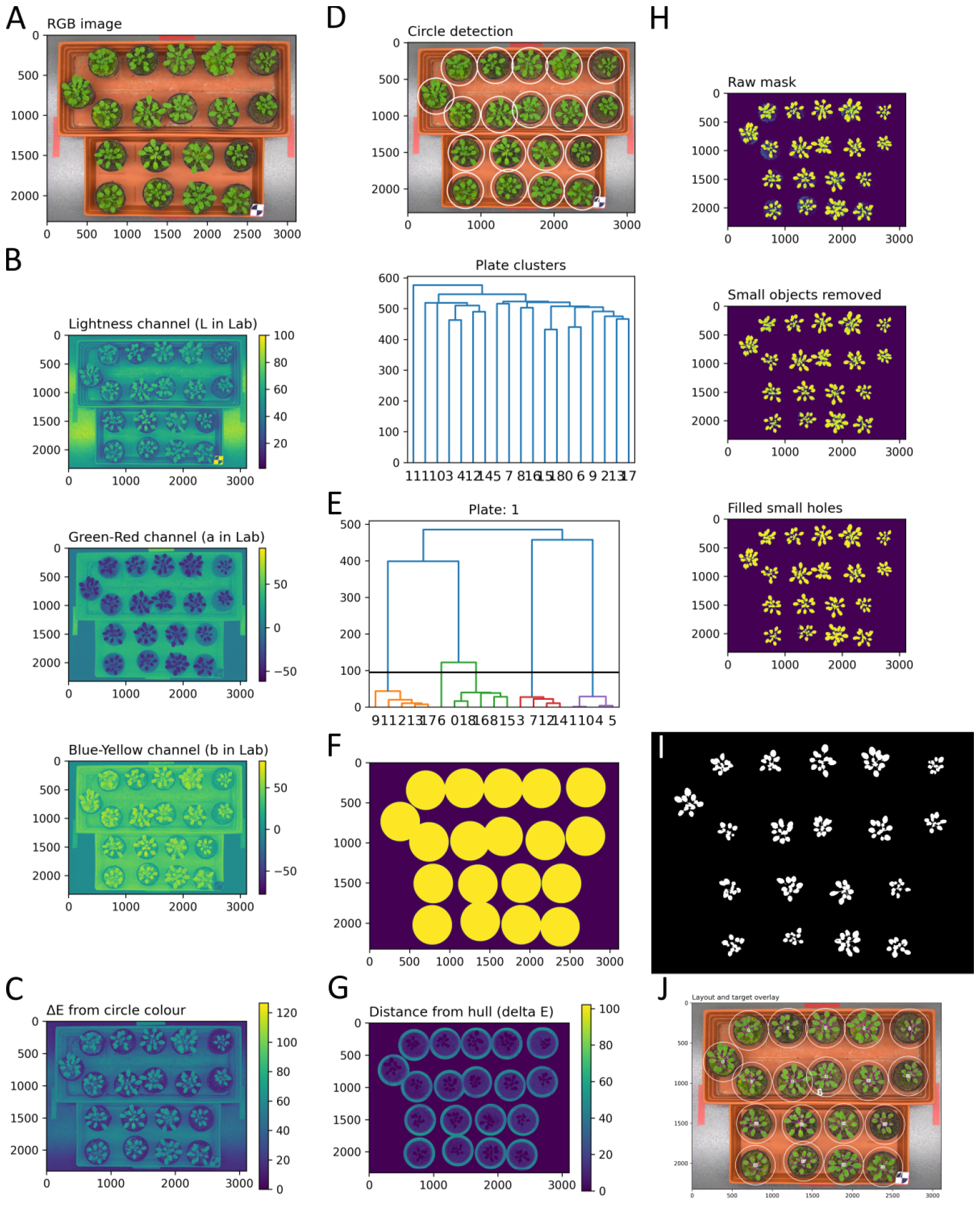
Representative image debugging output from area analysis on Arabidopsis. using the Ara2012 tray dataset obtained from http://www.plant-phenotyping.org/datasets (Minervini et al., 2015). A: RGB image. B: Image in each channel of CIELAB. C: Distance image from circle color. D: Circle detection overlay and plate clustering dendrogram. E: Row clustering for indexing including cut height. F: Layout mask. G: Distance image from target hull. H: Mask processing, from the initial delta cut-off with remove and fill steps. I: Target mask. J: Annotated overlay with IDs for the plate (1) and each unit(1-19) as well as the outline of the target mask.

In growth rate analysis from the Ara2012 series, a poorly fit model was identified corresponding to a region of interest for unit 8 (Fig. 4A). Multiple issues contribute to this poor fit, including herbivory, overlapping regions of interest and a need to extend the target hull (Fig 4B). This demonstrates how outlier detection in growth models can contribute to the identification of experimental and/or segmentation issues and prompt corrective action or consideration during subsequent analysis.

**Figure 4.**
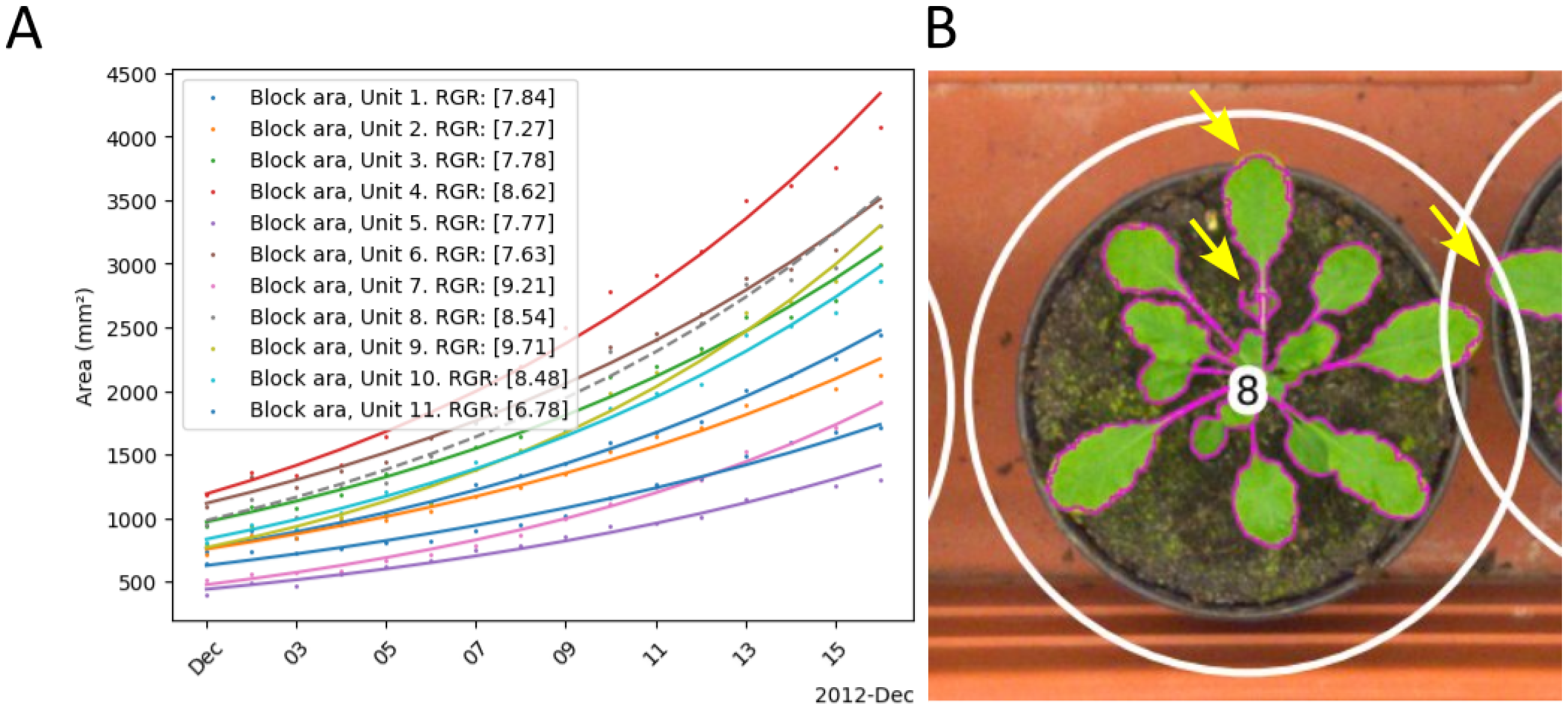
Relative growth rate analysis and the value of debugging images. using the Ara2012 tray dataset obtained from http://www.plant-phenotyping.org/datasets (Minervini et al., 2015). Image filenames were parsed to infer a block name and date for this analysis (see figure 3 and configuration in supplementary file “ara2012.conf”). A sample ID file was provided that separated units into groups, 1-11 as Tray 1 and 12-19 as Tray 2 (supplementary file “samples_ara2012.csv”). A: For each group, a plot is generated to display the straight line fit to log(area) over time, transformed back to units of area. The dashed line for unit 8 indicates the ModelFitOutlier flag was applied, suggesting a poor fit to the log-growth model and prompting further investigation. B: Inspection of overlay images identifies multiple issues indicated by arrows; vegetative damage that resembles herbivory by fungus gnat larvae, overlapping regions of interest and a need to extend the hull to include colors of pixels representing leaf tips, particularly where they overlap the lighter colored tray. Configuration for this figure is described in supplementary file “ara2012.conf” and the corresponding samples description is available as supplementary file “samples_ara2012.csv”.

In our usage with 8MP images, layout detection typically takes 10-30 seconds and image segmentation takes <5 seconds. A few additional seconds are required to load image data and generate debugging images. Area analysis provides an output file containing, for each target region; pixel count, scaled target area, median color of the target in RGB and CIELAB color spaces. Such color information has previously been used in other phenotyping applications such as detecting nutrient deficiency (Wiwart et al., 2009).

## DISCUSSION

Automated annotation of multiplexed images has previously been implemented in the PlantCV framework, relying on subject detection and assuming a true grid (plantcv.roi.auto grid) (Schuhl et al., 2022). Although the reported strategy is robust to some missing subjects, in AlGrow we have preferred to detect internal references to ensure consistent annotation. To support more flexible arrangements, detected circles are clustered into rows or columns rather than a true grid and multiple groups of these arrangements may also be included within a single image.

The visualization of color distributions in AlGrow facilitates the definition of a suitable margin for color based classification and can also help to improve experimental design through an intuitive representation of the underlying segmentation process. Together, the results from public datasets have highlighted the robustness of the hull in target color specification and the value of image debugging output and outlier detection during growth rate analysis (Figs. 2-4, S5 and Table S1). Although not demonstrated here, we expect AlGrow is also suitable in ecological applications that have previously applied alpha-hulls to the estimation of color volumes (Gruson, 2020).

AlGrow is suitable as a stand-alone application to perform area and growth rate analyses in plant and macroalgal phenotyping. AlGrow applies novel and effective methods for image segmentation and annotation, and provides thorough debugging and quality control output for parameter tuning and investigation of new or aberrant images. The software is easy to use, free and released open-source with compiled binaries for most platforms, making it suitable for a broad community of users. In addition to this article, a detailed guide and video tutorial explaining the use of the software and its features are available on the project page, https://github.com/marcusmchale/algrow.

## MATERIALS AND METHODS

### AlGrow architecture

AlGrow is developed in Python (>=3.10), is released open source (BSD-3 clause license) and provides a convenient cross-platform compatible graphical user interface (GUI) for calibration and analysis, along with a command line interface (CLI) for high throughput analyses. Parameters can be configured using the GUI in a desktop environment then written to a configuration file to apply the same analyses on other datasets or in other computational environments (Weisburd, 2023).

Matplotlib is used to annotate images and generate figures, including; debugging and quality control figures and summary plots describing growth (Hunter, 2007). Scikit-image is used extensively for; image import, color space conversion, edge detection, circle detection, pixel counting, fill and remove functions (Van der Walt et al., 2014). Numpy is used for fast and convenient computation and model fitting from array data, with Pandas also used for its convenient data structures and utilities (McKinney, 2010; Harris et al., 2020).

Configuration parameters may also be supplied as arguments to the CLI and are referenced in full form in this section, i.e. “--argument”. A launch option is available to remove noise with a bilateral filter (--denoise). Similarly, a downscale launch argument is provided (--downscale) that will reduce the image size during loading, though this currently has an interacting impact on all pixel distance parameters.

### Automated annotation

Distance (ΔE) from the chosen circle color (--circle_colour, Fig. 2A) is used to generate a gray-scale image for Canny edge detection (Fig. 2B) and Hough circle detection (Fig. 2A) in a specified range of sizes (--circle_diameter, --circle_variability) (Yonghong Xie and Qiang Ji, 2002). The region of interest for each detected circle can then be expanded or contracted (--circle_expansion). 238

Circles that are not completely within the image range (prior to expansion) are removed, then agglomerative clustering of circle center distances is performed to identify clusters of known sizes, termed plates (--circles_per_plate) (Müllner, 2011; Virtanen et al., 2020). The dendrogram cut height for this purpose is calculated from the expected distance between circle centers within a plate (--circle_separation, –circle_separation_tolerance and --circle_diameter). Plates are then clustered into rows or columns (--plates_cols_first) using their center position along the corresponding image axis. Plate rows or columns are separated with a cut-height of half the plate width (--plate_width). Circle ID incrementation is first determined by plate order, left to right and top to bottom, with options provided to reverse these (--plates_right_left/--plates_bottom_top).

Within plates, a grid-like arrangement of circles is assumed to perform a rotation correction, enforcing alignment with the image axes to ensure consistent annotation order. The angle of rotation is determined by examining the two angles on each of four corners relative to their most closely aligned image axis, with the median of all eight angles selected. Rotated positions are then used for agglomerative clustering into rows or columns (–circles_cols_first) along the corresponding image axis. Circle rows or columns are separated with a cut height of one quarter of the circle diameter (--circle_diameter). As for plates, the default incrementation order is left to right and top to bottom and options are provided to invert these (--circles_right_left/--circles_bottom_top).

Where absolute positions do not change, a fixed layout may be detected on one image, saved and later loaded (--fixed_layout) to avoid repeating these calculations. Layout specification may also be ignored to process whole images, though it is recommended to start with this step to mask background where possible before proceeding to target hull segmentation.

### Segmentation

We employed the Open3D library extensively in GUI development and hull distance calculations (Zhou et al., 2018). Pixel colors are down-sampled to voxels (--voxel_size) and the signed distance to the hull surface is computed for each voxel by ray-casting. In the GUI, a mapping of pixel to voxel is used to highlight pixels whose color is within the hull or within ΔE (--delta) of its surface. However, during area analysis the distance is computed for each pixel.

Convex and alpha hull construction is achieved in using the Trimesh and Alpha Shape Toolbox libraries, respectively (Bellock, 2015; Dawson-Haggerty et al., 2019). When the alpha parameter is set to zero, the convex hull is used. For other values of alpha, the specified value is the maximum distance (ΔE) between two points for these to be considered as edges in hull construction.

In creating a hull from a supplied image mask, voxels mapped to pixels that are located within the foreground mask are first filtered by the supplied min pixels. A hull is then constructed from these at the current alpha value and vertices of this are retained as the current voxel selection. Due to computational demand, it is not recommended to combine low values of alpha (≲5) with low values of min pixels (≲5) except on small (≲1 megapixel) images.

### Growth analysis

Multiple image files may be selected for segmentation with the current hull and delta using concurrent processing (--processes). A regular expression can be provided to parse filenames to determine date, time and optionally a block for each image. These details are then included in the output area file and used in subsequent relative growth rate (RGR) analysis. 288

RGR is calculated as the slope of a line of best fit to log transformed area over time. Plots of area over time, with the line fit in log(area) transformed back to units of area and box-plots of RGR per sample group (Fig. 5), are all generated according to labels provided in a mapping file (--samples). This mapping file can be generated as a “.csv” file using spreadsheet software. To assist in the identification of any experimental issues, the “ModelFitOutlier” flag is applied to fits where the residual sum of squares (RSS) is higher than the median RSS across all fits plus 1.5 times the inter-quartile range.

### Arabidopsis image analysis

In Ara2012 images the individual trays could not be distinguished as the assumption of circles within trays being closer together than circles between trays is not met. In Ara2013 images, the last image failed layout detection due to significant growth over the margins in one pot. As such, for consistency, the first and second last image were used for hull training.

## FIGURE LEGENDS

**Figure S1: Scale definition in the AlGrow GUI**. A line may be drawn across a span of known dimensions at the relevant depth of field, here the width of the culture plate, and the measured pixel distance is displayed in the text box (px). The known distance in millimeters (mm) is entered and a scale (px/mm) is calculated.

**Figure S2. Edge detection is performed on a distance image**, reflecting the ΔE from a specified circle color (--circle_colour). A: In the AlGrow GUI, a loaded image (here Palmaria palmata in the macroalgal phenotyping apparatus) is displayed (right) and pixel colors may be selected. B: Selected colors are displayed as buttons (bottom left) with labels corresponding to CIELAB coordinates. Each selected color may be removed by clicking on the corresponding button, or all selected colors may be removed by clicking “Clear”. The median of these selected color values in CIE LAB, the current “circle colour”, is displayed as a button below “Clear” and used to prepare the distance image for Canny edge detection (displayed at right).

**Figure S3. Failed layout detection prompts an error message**, here an attempt to detect 7 plates with only 6 circles per plate raises the insufficient plates detected error. When an error occurs, an overlay of the image with detected circles alongside the dendrogram of circle center distances is displayed. The cut-height (black line) is included in the dendrogram and users can consider this figure to adjust any necessary parameters.

**Figure S4. A cuboid selection with boundaries determined from an interactive session using ImageJ** is presented to depict thresholds and demonstrate their limitations for this image (Dice = 0.70). In particular, the low exposed regions cannot be captured within a single cubic selection without selecting other background. Alternate viewing angles are also presented (inset) to more clearly represent this issue. Configuration loaded into AlGrow for this figure is described in supplementary file “palmaria_cube_imagej.conf”.

**Figure S5. An alpha hull target specification with fill and remove** settings applied used to create an accurate image mask. The Dice coefficient with the mask generated using this configuration is approximately but not exactly 1.0 due to voxel down-sampling applied in the GUI but not during formal area analysis. Alternate viewing angles are presented (inset) to best depict the concave surfaces of this hull. Configuration loaded into AlGrow for this figure is described in supplementary file “palmaria_alpha.conf”.

## Supplementary files

Tables:

- **Table S1: Dice coefficient, accuracy, sensitivity and specificity** for the analysis of the Ara2012 and Ara2013 tray datasets performed using the configuration defined in supplementary files “ara2012.conf” and “ara2013.conf”. 413

Images:

- palmaria.jpg
- palmaria_alpha_mask.png
- Configuration files:
- ara2012.conf
- ara2013.conf
- palmaria.conf
- palmaria_alpha.conf
- palmaria_cube_imagej.conf

## ACKNOWLEDGMENTS

We thank Matthias Schmid, Serena Rosignoli and Gaelle Bihan for images of *Palmaria palmata*. We also thank Mathias Schmid for the suggestion to report typical color for each target.

